# Unbiased integration of single cell multi-omics data

**DOI:** 10.1101/2020.12.11.422014

**Authors:** Jinzhuang Dou, Shaoheng Liang, Vakul Mohanty, Xuesen Cheng, Sangbae Kim, Jongsu Choi, Yumei Li, Katayoun Rezvani, Rui Chen, Ken Chen

## Abstract

Acquiring accurate single-cell multiomics profiles often requires performing unbiased *in silico* integration of data matrices generated by different single-cell technologies from the same biological sample. However, both the rows and the columns can represent different entities in different data matrices, making such integration a computational challenge that has only been solved approximately by existing approaches. Here, we present bindSC, a single-cell data integration tool that realizes simultaneous alignment of the rows and the columns between data matrices without making approximations. Using datasets produced by multiomics technologies as gold standard, we show that bindSC generates accurate multimodal co-embeddings that are substantially more accurate than those generated by existing approaches. Particularly, bindSC effectively integrated single cell RNA sequencing (scRNA-seq) and single cell chromatin accessibility sequencing (scATAC-seq) data towards discovering key regulatory elements in cancer cell-lines and mouse cells. It achieved accurate integration of both common and rare cell types (<0.25% abundance) in a novel mouse retina cell atlas generated using the 10x Genomics Multiome ATAC+RNA kit. Further, it achieves unbiased integration of scRNA-seq and 10x Visium spatial transcriptomics data derived from mouse brain cortex samples. Lastly, it demonstrated efficacy in delineating immune cell types via integrating single-cell RNA and protein data. Thus, bindSC, available at https://github.com/KChen-lab/bindSC, can be applied in a broad variety of context to accelerate discovery of complex cellular and biological identities and associated molecular underpinnings in diseases and developing organisms.

## Introductions

Advances in high-throughput single-cell technology such as single-cell RNA-sequencing (scRNA-seq) ^1^ and mass cytometry ^2^ have enabled systematic delineation of cell types based on thousands to millions of cells sampled from developing organisms or patient biopsies. For example, recent application of combinatorial indexing based technology has generated the transcriptomic and chromatin accessibility profiles of millions of cells in developing human fetus samples ^3^. Rare cell types and complex cellular states, however, remain challenging to discover, which necessitates the development of multiomics technologies to simultaneously measure other cellular features, including DNA methylation ^4,5^, chromatin accessibility ^6-8^ and spatial positions ^9,10^ in the same cells. Although available single-cell multiomics technologies ^8,11-14^ can profile thousands to millions of cells per experiment, the cost of the experiments is still quite high ^15^; and the data generated are often of lower throughput than those generated by unimodal technologies. These restrictions necessitate the development of computational approaches that can accurately integrate multiple data matrices generated by different technologies from the same biological samples to acquire an accurate characterization of cellular identity and function.

However, different technologies create data matrices of different rows and columns, which correspond to different sets of cells and different types of features. How to align cells and features simultaneously across matrices is a core computational challenge. When the two sets of cells are sampled uniformly from the same biological sample, it is safe to assume that there exists an optimal way to align together cells of similar identities and features associated with these identities. This is mathematically challenging, however, as there are many possible ways to simultaneously align a large number of cells and features. To address this challenge, existing computational approaches followed two directions: 1) aligning features empirically before aligning cells ^16-19^; 2) obtaining separate embeddings for each modality, followed by performing unsupervised manifold alignment ^20-22^. Taking integration of scRNA-seq and singe cell assay for transposase accessible chromatin sequencing (scATAC-seq) as an example, the first category of methods require constructing a “gene activity matrix” from scATAC-seq data by counting DNA reads aligned near and within each gene ^23^. This strategy considers only the basic cis-regulatory relations and ignores long-range, trans-regulatory relationship established via other regulatory elements such as enhancers ^6^, which are often critical to decipher cell identities. It also substantially simplifies (or loses) multifactorial relations between transcription factors (TF) and target genes ^24^ Based on pre-aligned features generated by such empirical rules, Seurat applies canonical correlation analysis (CCA) and mutual nearest neighbors (MNNs) to identify cells anchoring the two data matrices ^17^; LIGER uses an integrative non-negative matrix factorization (iNMF) to delineate shared and dataset-specific features ^19^; Harmony projects cells onto a shared embedding using principle components analysis (PCA) and removes batch effects iteratively ^18^. All these programs suffer from the aforementioned limitations and thereby cannot yield a comprehensive, unbiased gene regulatory network, particularly when chromatin changes are asynchronous from RNA transcriptions in cells undergoing state transitions ^25^. The second category of methods ^20-22^ do not require prior feature alignment and are fully unsupervised. However, they depend heavily on the assumption that feature variation across cells is driven by a few latent variables in both modalities ^22^. This assumption can get violated easily in datasets of complex biology involving dynamic processes such as differentiation, reprogramming and transdifferentiation ^22^.

In this study, we develop a novel computational tool called bindSC (bi-order integration of single-cell data). The key algorithm implemented in bindSC is called bi-CCA (bi-order canonical correlation analysis). Bi-CCA learns the optimal alignment among rows and columns from two data matrices generated by two different experiments. The alignment matrix derived from bi-CCA can thereby be utilized to derive *in silico* multiomics profiles from aligned cells.

We assess our method on several challenging multimodality integration tasks between 1) transcriptomic and chromatin accessibility data, 2) transcriptomic and spatial transcriptomic data, and 3) transcriptomic and proteomic data. We validate scRNA-seq and scATAC-seq integration accuracy using datasets obtained directly from multiomics technologies, including a novel mouse retina cell atlas created by the 10x Genomics Multiome ATAC+RNA kit. We show that bindSC enables comprehensive characterization of epigenetic regulatory states in a lung adenocarcinoma cell-line A549 in response to dexamethasone treatment. And bindSC can align mouse retina cell types accurately, for multi-subtype bipolar cells and rare horizontal cells. Moreover, bindSC enables unbiased integration of spatial transcriptomics data with scRNA-seq data on mouse brain cortex samples, as well as single-cell RNA data with protein data from peripheral blood mononuclear cells. BindSC is implemented as an open-source R package available at https://github.com/KChen-lab/bindSC.

## Results

### Bi-order integration of multi-omics data

BindSC takes as input two single-cell data matrices (**X** and **Y**) generated uniformly from the same cell population by two different technologies (**Fig. 1a**). In most single-cell multi-omics integration tasks, neither the alignment between the cells in **X** and those in **Y**, nor the alignment between the features in **X** and those in **Y** is known. BindSC employs a bi-CCA algorithm developed in this study to address this challenge (**Fig. 1b**). Briefly, bi-CCA introduces a gene score matrix **Z** to link **X** and **Y**. The gene score matrix has the same rows as does **X** and the same columns as does **Y**. To reduce computational cost, **Z** can be initialized based on prior knowledge. Taking integration of scRNA-seq and scATAC-seq as an example, the gene score matrix can be initialized using the “gene activity matrix” estimated by other programs such as Seurat. Bi-CCA then iteratively updates **Z** to find an optimal solution which maximizes the correlation between **X** and **Z** and between **Y** and **Z** in the latent space simultaneously. Details about this iterative procedure can be found in **Methods** and **Supplementary Fig. 1a**.

**Fig. 1.**
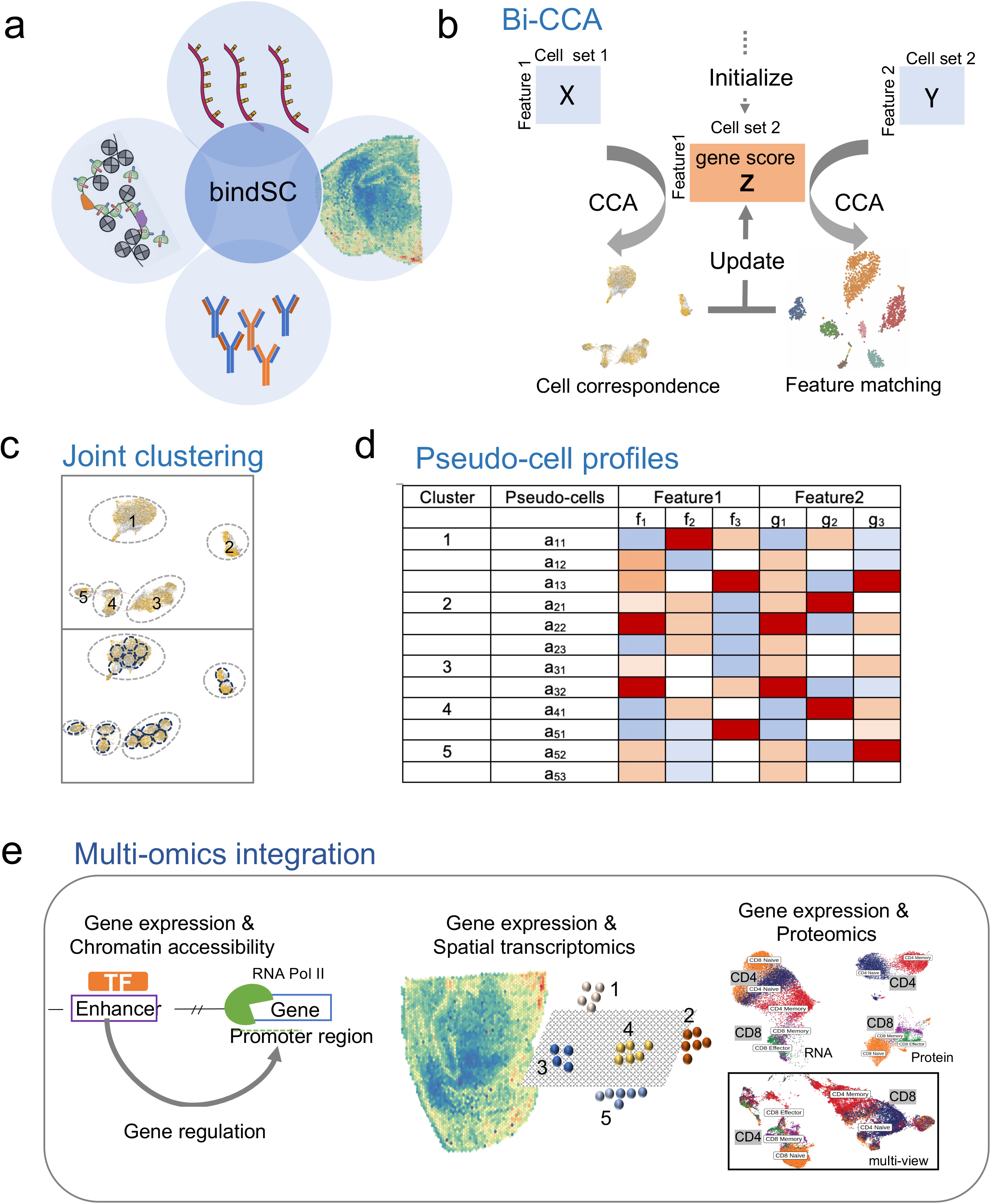
Overview of bindSC. BindSC takes as input two data matrices produced by different modalities from the same cell population (**a**). The modalities may include transcriptomes, epigenomes, spatial transcriptomes and proteomes. Bi-order integration of two modalities (**X** and **Y**) with unpaired cells and unmatched features using Bi-CCA algorithm **(b)**. In the data matrices, each row represents one gene/locus, and each column represents one cell. The gene score matrix **Z** that links the first modality with the second one is initialized by prior gene activity modeling (see **Methods**). Bi-CCA algorithm aims to update gene score matrix **Z** iteratively by maximizing the correlation of between **X** and **Z** and between **Y** and **Z** simultaneously. Based on canonical correlation vectors (CCVs) in the derived latent space, K-nearest neighbor (KNNs) clustering is performed to define cell types in both modalities **(c)**. Within each cell-type cluster, KNN clustering is further performed at a higher resolution to define pseudo-cells consisting of 10s cells from both modalities. *In silico* multimodal profiles are constructed from cells assigned to the same pseudo-cell **(d)**. The color in each box indicates the relative level of each feature, with white corresponding to missing values. The multiomics feature profiles enable us to 1) link genes to regulatory elements, 2) map RNA expressions to spatial locations and 3) delineate cells by both RNA and protein signatures **(e)**.

Bi-CCA outputs canonical correlation vectors (CCVs), which project cells from two datasets onto a shared latent space (referring below as “co-embedding”). A K-nearest neighbor (KNN) graph is constructed based on Euclidean distances observed in the latent space, followed by modularity optimization techniques to partition the KNN into highly interconnected subgraphs, each of which corresponds to a putative cell type or state (**Fig. 1c**). Within each cluster, sub-clustering using similar strategies is further performed to derive what we call pseudo-cells (**Methods**). Each pseudo-cell encloses tens of cells from both datasets and thus has a consensus multiomic profile summarized from constituting cells (**Fig. 1c-d**). The joint multiomic profiles thus enable 1) characterizing gene and chromatin-accessibility relations from aligned scRNA-seq and scATAC-seq data; 2) associating transcriptomic profiles with spatial locations from aligned scRNA-seq and spatial transcriptomic data; 3) associating transcriptomic profiles with proteomic profiles from aligned scRNA-seq and CyTOF data, and so on (**Fig. 1e**).

### Benchmarking bindSC performance on simulation datasets

Existing integration methods such as Seurat, LIGER, and Harmony require pre-aligning features across modalities, i.e., compressing cell-peak matrices obtained from scATAC-seq onto cell-gene-activity matrices based on reference genome annotations. BindSC overcomes that restriction: its generic mathematical formulations allow free alignment amongst features to be established from data.

Under our formulation (**Methods**), **Z** has features (rows) aligned with **X** and cells (columns) aligned with **Y**. The introduction of **Z** enables bi-order alignment of the cells and the features, respectively.

To quantify how much this step matters to overall integration accuracy, we performed a set of simulation experiments. We started by creating a dataset **X** consisting of 3 cell clusters (types), each having 333 cells and 1,000 genes using Splatter ^26^ (**Supplementary Fig. 2a**). We created a second dataset **Y** and made it identical to **X**: **X** = **Y**. We then constructed a gene score matrix **Z** from **Y** by permuting a fraction of features (rows), termed misalignment rate (MR), into different orders. The features between **Z** and **Y** are perfectly aligned if MR equals 1 and are independent if MR equals 0. We further added white noise on all the entries of **Z** at a given signal-noise-ratio (SNR) level.

We then provide (**X**, **Z**) as input to the other methods (**Supplementary Fig. 2b**), mimicking how they perform integration, while provide both (**X**, **Z**) and (**Y, Z**) to bindSC (**Supplementary Fig. 2c**). As described, rather than taking **Z** as it is from the input, bindSC will iteratively update **Z** until reaching convergence.

Since we know the true cell type and dataset origin of the cells in these experiments, we can assess the integration performance in terms of cell type classification accuracy and dataset alignment accuracy in the co-embeddings. It is necessary to measure both types of accuracy, as a high cell type classification accuracy can be achieved by simply projecting cells onto local clusters without achieving uniform mixing of the two datasets. Similarly, a high dataset alignment accuracy can be achieved by uniformly mixing cells from the datasets, regardless of their cellular identity. We used Silhouette score for measure cell type classification accuracy and alignment mixing score to measure the dataset alignment accuracy (**Methods**). We compared bindSC, CCA, Seurat, LIGER and Harmony under default settings (**Supplementary Note 1**).

We obtained results from a range of MRs under SNR = 0.25 (**Fig. 2**). When there was no feature misalignment (MR = 0), all methods achieved good performance. Even under this ideal scenario, bindSC achieved the highest Silhouette score (> 0.75) (**Fig. 2a**). The worse performance of other methods can be explained by the noise introduced to distort the manifold structures between **X** and **Z**. CCA showed better performance than Seurat, which may be due partly to label transferring errors introduced by Seurat’s empirical anchor-based alignment approach. As MR increased from 0 to 0.9, the Silhouette score for bindSC remained stable (> 0.7), while all the other methods showed a decreasing trend, especially for LIGER and Harmony. Harmony worked well when MR ≤ 0.15 (**Fig. 2a-b**) but had a substantial drop on Silhouette score (< 0.1) when MR > 0.15. In addition, its alignment mixing score dropped to 0 when MR > 0.2, with no mixing of cells from **X** and **Z** in the co-embedding UMAP (**Fig. 2b**; MR = 0.5). Harmony takes cell coordinates from a reduced dimensional PCA space and runs an iterative algorithm to adjust for dataset-specific effects. When MR > 0.15, cells from **X** and **Z** already formed two dis-joint groups, which made the downstream integration impossible for Harmony. The Silhouette score of LIGER showed fluctuations but was always lower than 0.4. LIGER utilizes an integrated nonnegative matrix factorization (iNMF) method to identify shared and dataset-specific metagenes across two datasets. If it worked as designed, the errors caused by feature misalignment should be contained within dataset-specific modules. However, variance explained by the data-specific modules appeared to be small (< 1%). When MR ≥ 0.95, all methods including bindSC failed to achieve reasonable integration. That was expected as **X** and **Z** (as well as **Y** and **Z**) became nearly independent.

**Fig. 2.**
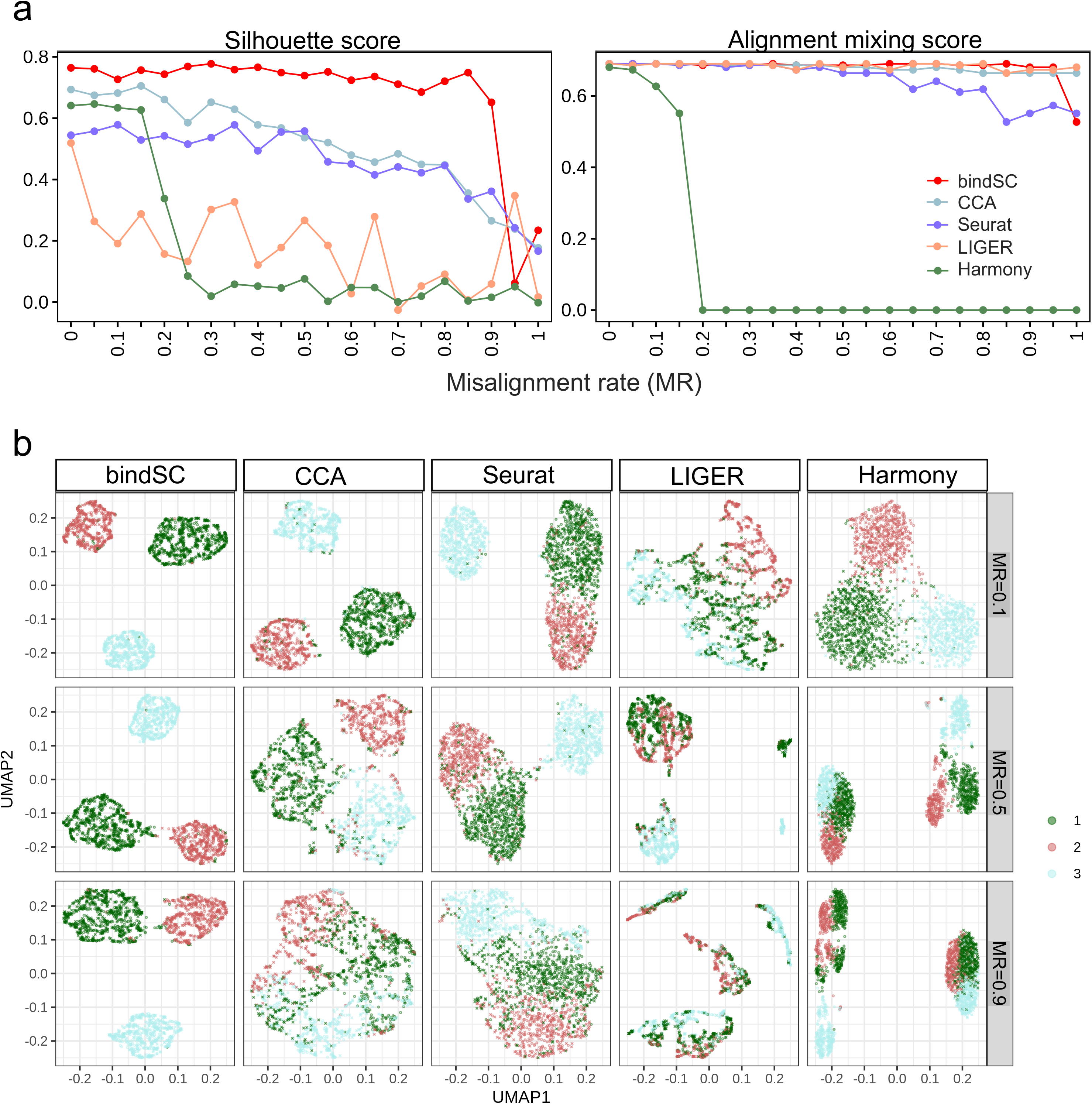
Benchmarking bindSC performance on simulation datasets. Comparison of bindSC to CCA, Seurat, LIGER, and Harmony based on Silhouette score and alignment mixing score **(a)**. The dataset contains 1,000 genes and 1,000 cells equally distributed in 3 cell types. Signal-to-noise ratio (SNR) was set at 0.25. X-axes denote the misalignment rates (MR) between features in the two datasets, which ranges from 0 to 1. The features between two datasets have perfect match if MR = 0 and are unrelated if MR = 1. UMAP views of the co-embeddings generated by bindSC, CCA, Seurat, LIGER, and Harmony **(b)**. From top to bottom are results with MR = 0.1, 0.5, and 0.9, respectively. Each point denotes one cell that is colored based on its true cell type label (red, green, or cyan).

As expected, increasing SNR level worsened the integration performance for most of the methods except bindSC. For example, both CCA and Seurat had acceptable performance under MR = 0.5 and SNR = 0 (**Supplementary Fig. 3a**), but Seurat failed to separate cell type 2 and 3 accurately when SNR = 0.25 or 0.5 (**Supplemental Fig. 4a**; **Fig. 2**). For SNR = 0.5, Harmony failed in both alignment mixing (< 0.2) and classification (= 0) accuracy, even when MR was as low as 0.1 (**Supplementary Fig. 4**).

We repeated the above experiments by increasing the number of cells to 5,000 and 10,000, respectively. Similarly, bindSC showed robust performance regardless of MR and SNR levels, which was not achieved by other methods (**Supplementary Tables S2-3**). Overall, the simulation results demonstrated that bindSC is robust to bias introduced by noise in the data and via pre-aligning features, thanks to its ability to align both cells and features simultaneously.

### Integrating single cell epigenomic data with single cell transcriptomic data

Integrating single cell epigenomic data with single cell transcriptomic data obtained from unimodal technologies provides an opportunity to decipher epigenetic regulatory mechanisms underpinning cell transcriptomic identity. We examined the performance of bindSC in integrating the scRNA-seq and scATAC-seq data derived from lung adenocarcinoma (A549) cells after 0, 1, and 3 hours of dexamethasone (DEX) treatment ^6^. This dataset was generated using a combinatorial indexing-based coassay (sci-CAR), which enabled jointly measurement of chromatin accessibility and transcriptome in the same cells. In this dataset, 6,005 cells have sci-RNA-seq profiles and 3,628 cells have sci-ATAC-seq profiles. Among them, 1,429 cells have both RNA-seq and ATAC-seq profiles, which can be used as a gold standard for evaluating integration accuracy of various methods (**Methods**).

For comparison, we ran the 4 methods on the same data and derived *in silico* co-embeddings. There was relatively clear separation between cells acquired at 0 hour and those at 1 or 3 hours in the co-embeddings (**Fig. 3a**). In terms of classifying cells by time, bindSC achieved the highest Silhouette score and Harmony the second, whereas Seurat had the lowest score with many sub-clusters in its co-embedding (**Fig. 3a-b**). As to alignment accuracy, bindSC and Harmony had similar scores, whereas Seurat received a relatively low score (**Fig. 3b**). Similar trends were observed in a previous study analyzing the same dataset ^27^. As suggested by simulation, the low alignment mixing score of Seurat was likely attributable to bias introduced in its anchor-based integration.

**Fig. 3.**
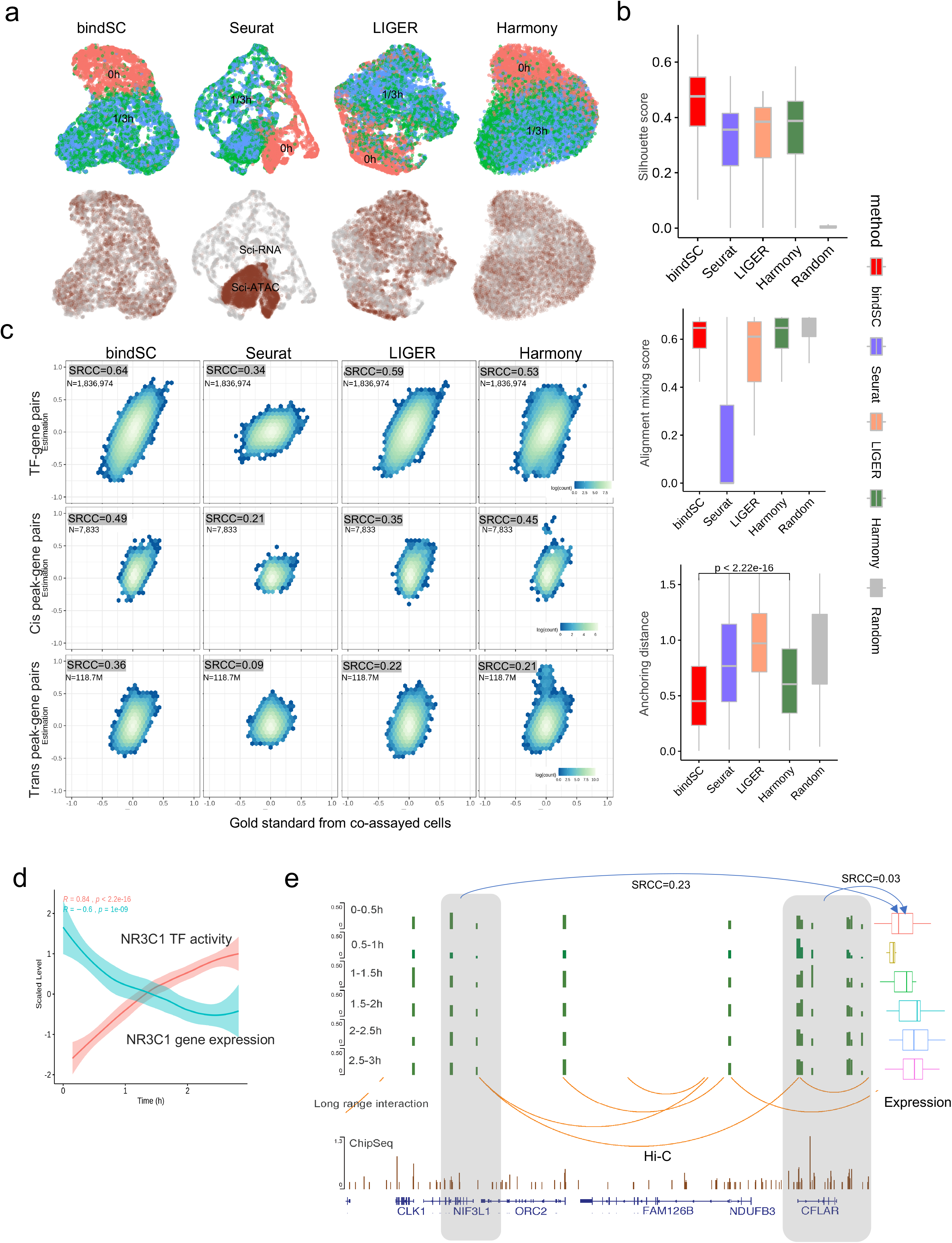
Benchmarking bindSC performance on the DEX-treated A549 cell-line data. UMAP of cells from DEX-treated A549 cell-line data for bindSC, Seurat, LIGER and Harmony respectively, colored by collection time (red: 0 hour, green: 1 hour and blue: 3 hour) on the top panel and by technologies (grey: sci-RNA and brown: sci-ATAC) on the bottom panel **(a)**. Comparison of the 4 methods based on Silhouette score (top), alignment mixing score (middle) and anchoring distance (bottom) **(b)**. Also included for comparison are metrics from randomly selected cells. Accuracy of *in silico* multiomics profile in pseudo-cells **(c)**. The TF-gene correlation is quantified in each pseudo-cell (top panel) by calculating for each TF-gene pair (1.8 million pairs total) a Spearman’s rank-order correlation coefficient (SRCC) between the TF activity level, inferred based on motif enrichment in the ATAC data, and the gene expression level in the RNA data. The peak-gene correlation is quantified by calculating for each peak-gene pair a Spearman’s rank-order correlation coefficients (SRCC) between a normalized ATAC peak level and a gene expression levels for the cis-peaks (middle panel, 7,833 pairs) and the trans-peaks (bottom panel, 118.7 million pairs), respectively. X-axes are the SRCCs estimated from the co-assayed cells, which serve as the gold standard, while Y-axes are the SRCCs estimated from the pseudo-cells generated by each method. The overall concordance between X and Y are further quantified using a single SRCC shown on the up-left corner of each subfigure. Cis is defined as gene bodies plus 2,000 bps upstream. Reconstructing the gene expression and the TF activity level (Y-axes) of *NR3C1* using bindSC pseudo-cells **(d)**. X-axis is the averaged treatment time of the cells in each pseudo-cell. A genome browser view showing putative regulatory relations between an accessible distal site chr2:201770437-201770992 and the gene *CFLAR* **(e)**. The 6 tracks at the top show ATAC peak levels and gene expression levels at six time points. The track in the middle shows chromatin interactome from published Hi-C data. The bottom track shows the *NR3C1* binding targets (ChIP-Seq) peaks published in an independent study ^28^.

A perfect integration method would place the two instances of the 1,429 co-assayed cells onto identical locations in the co-embeddings. We leveraged this expectation to compare the accuracy of various methods. We defined a metric, called anchoring distance that measures the normalized Euclidean distance between the two instances of a co-assayed cell in the co-embeddings (**Methods**). BindSC achieved substantially shorter anchoring distances than the other methods (*p* < 2.2e-16; Student t-test; **Fig. 3b**).

We further compared how accurately TF (or peak) -gene correlations can be inferred from the co-embeddings produced by each method. For a fair comparison, we applied the same bindSC workflow to derive pseudo-cells for the 4 methods (**Methods; Supplementary Note 2**).

For each TF-gene (and peak-gene) pair, we calculated a Spearman rank correlation coefficient (SRCC) between the TF activity (and normalized peak) level and the gene expression level in the pseudo-cells (**Methods**). We repeated the same calculation in the co-assayed cells to create a gold standard. For each of the 4 methods in 3 types of relations: TF-gene, cis-peak-gene and tran-speak-gene, we calculated a summary SRCC between the SRCCs obtained from the pseudo-cells and the SRCCs obtained from the co-assayed cells. The summary SRCCs resulting from bindSC were consistently higher than those obtained from Seurat, LIGER and Harmony in all the categories of comparison, indicating that the bindSC multiomic profile had the highest accuracy.

We further examined the peak-gene association identified from the co-assayed cell profiles and found 585 trans-peak-gene pairs being supported by isogenic Hi-C data generated in an independent study ^28^. Compared with other approaches, bindSC derived peak-gene SRCCs of the highest level of agreement with those observed in the co-assayed cells (**Supplementary Fig. 5**). Among the 585 trans-peaks, 470 appeared more strongly correlated with the corresponding gene expression levels than did the corresponding cis-peaks. One example was the gene *CFLAR* and a trans-peak at chr2:201,770,437-201,770,992, which is 200-kb upstream of *CFLAR* transcription start site, spanning over three genes (**Fig. 3e**). The SRCC of this pair was 0.32 in the co-assayed cells. It was lower but comparable (0.23) in the bindSC pseudo-cells, however, became substantially lower (< 0.11) in the pseudo-cells generated by the other methods (**Supplementary Fig. 5**).

The DEX treatment specifically targets the glucocorticoid receptor encoded by *NR3C1,* a TF that activates the mRNA transcription of a handful of downstream genes. BindSC accurately reconstructed the gene expression and TF activity kinetics of *NR3C1* (**Fig. 3d**), consistent with what was depicted in the original study ^6^ using the co-assayed cells: the *NR3C1* expression level decreased over time while its activity level increased; Even the slowing down trend of *NR3C1* activity was captured.

We further evaluated the performance of bindSC in integrating scRNA-seq and scATAC-seq on another available multi-omics dataset generated recently by SHARE-seq technology^14^. There were a total of 37,774 cells from mouse skin tissues that had paired RNA and ATAC profiles. Compared with other methods, bindSC again achieved significantly shorter anchoring distances (**Supplementary Fig. 7**; **Supplementary Note 4**).

### Comprehensive evaluation using a novel mouse retinal cell atlas

For comprehensive evaluation and comparison, we generated a novel multi-omics dataset from single nuclei of wild type mouse retina. Mouse retina is heterogeneous, composed of multiple neuronal and non-neuronal cell types, including five major neuron classes: photoreceptors (rods and cones), retinal ganglion cells (RGC), horizontal cells (HC), bipolar cells (BC), amacrine cells (AC), and a non-neuronal Müller glial cell (MG) ^4,29,30^. While we ^29^ and others ^31-33^ have provided high-resolution single cell transcriptomic profiles of whole retina or specially sorted cell types on mouse and human retina tissue, little is known on the single-cell chromatin landscape of mouse retina tissue. Numerous studies ^34-36^ demonstrate the importance of transcription factors (TFs) on establishing or maintaining the chromatin landscapes that define retina cell identity. Therefore, integration of ATAC and RNA profiles at single cell resolution provides an exciting opportunity to comprehensively characterize cell types and rare cell subtypes in mouse retina.

We applied the newly released 10x Genomics Multiome ATAC+RNA kit on nuclei suspension acquired from adult mice retina samples. After performing standard quality control, we obtained an atlas of 9,383 nuclei of high-quality ATAC+RNA profiles. To define cell types, we first clustered the RNA and the ATAC data individually. Nineteen (19) clusters were identified from the RNA data alone, which included all the known major cell types with some subtypes identified: rod, BC (BC1~BC10), AC, RGC, cone, HC, MG and retina progenitor cells (RPC) (**Fig 4a and Supplementary Fig. 8**). Nineteen (19) clusters were also identified from the peak files of the ATAC data alone (**Fig. 4b**). Although known cell types appeared to be well separated in both modalities, there were some noticeable differences. For example, RGC cells and rod cells were separated clearly in the RNA data but partly blended together in the ATAC data, whereas ACs and RGC cells were blended in the RNA data but well separated in the ATAC data. Interestingly, all the 10 BC cell subtypes, defined based on RNA expression levels, were well separated in the ATAC data except for BC1 and BC6. However, after reducing ATAC data to gene level in a gene activity matrix, the cell types became considerably harder to delineate (**Fig. 4c**).

**Fig. 4.**
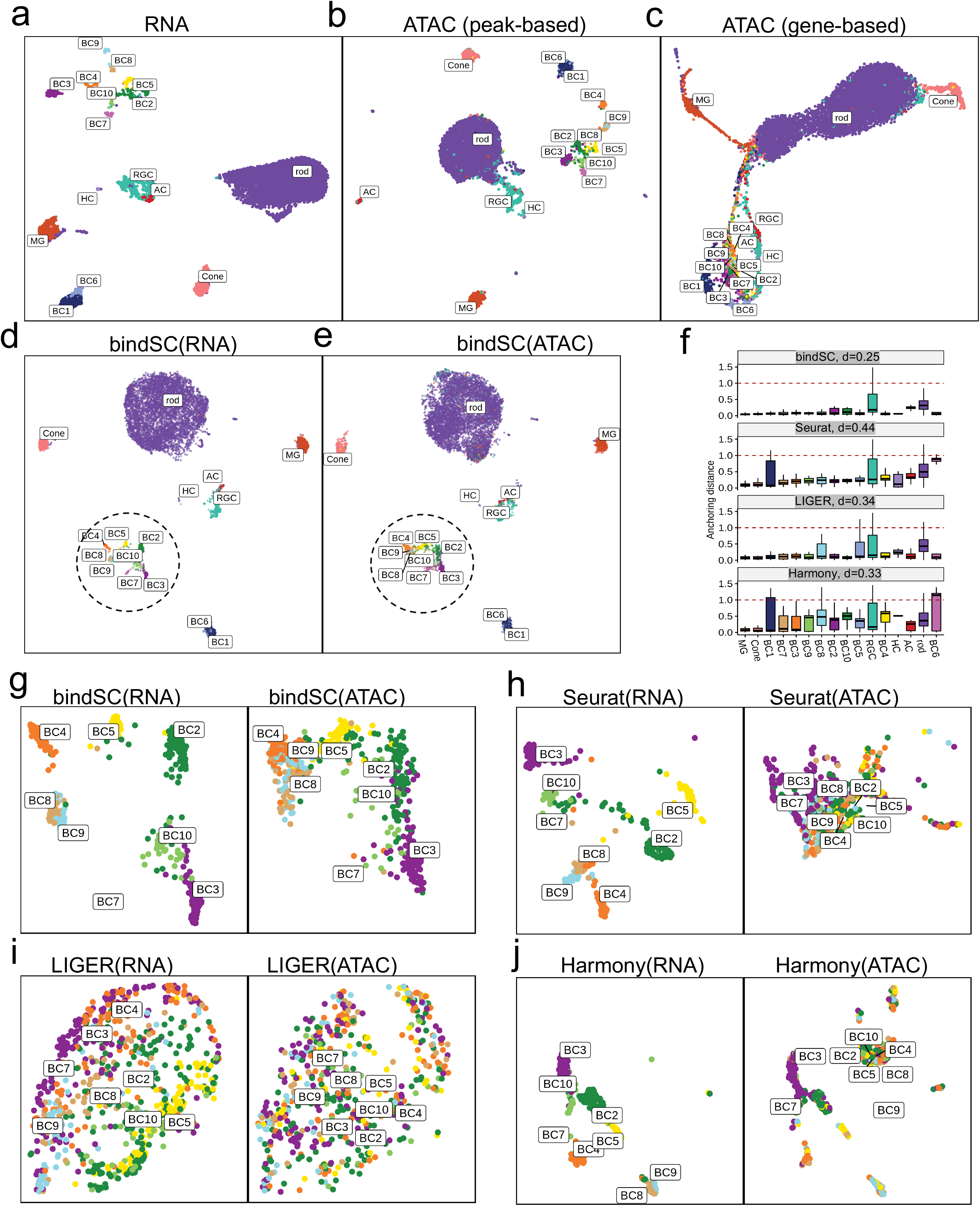
Integrating single-cell RNA-seq and ATAC-seq on a mouse retinal cell atlas. UMAP views of 9,383 mouse retina cells based on gene expression levels in the RNA-seq data **(a)**, chromatin accessibility peak profiles in the ATAC-seq data **(b)**, gene-level collapsed chromatin accessibility profiles **(c)**. The cells are colored by cell types annotated based on RNA expression levels (**Supplementary Fig. 8**). BindSC co-embeddings for the cells in the RNA-seq data **(d)** and those in the ATAC-seq data **(e)**. Anchoring distances resulting from bindSC, Seurat, LIGER and Harmony **(f)**. The median anchoring distance for each cell type was highlighted as a bold horizonal bar in the middle of each box in each panel. The dotted line denotes the anchoring distance based on random guesses. Zoomed out UMAP views for the BC cells in the co-embeddings generated by bindSC (**g**), Seurat (**h**), LIGER (**i**), and Harmony (**j**). Integration results for all the cell types can be seen in **Supplementary Fig. 9**. RGC: retinal ganglion cells; HC: horizontal cells; BC: bipolar cells; AC: amacrine cells; MG: Müller glial cell.

To obtain *in silico* multiomics profiles, we ran bindSC together with three other methods on the data without using the known cell correspondence. As shown in the co-embedding UMAP (**Fig. 4d-e**), bindSC successfully aligned cell types across modalities, with most cell types well separated out (**Fig. 4d-f**). Interestingly, bindSC successfully aligned the HCs, which is quite rare in the dataset (23 cells, <0.25% abundance). None of the other methods aligned the HCs correctly as it was already difficult to separate the HCs from the ACs in the gene-level chromatin profiles (**Fig. 4c and Supplementary Fig. 9**), the input to the other methods. Overall, the anchoring distances in the co-embeddings generated by bindSC were considerably smaller than those generated by the other methods in all the cell types assessed (**Fig. 4f**).

Note that bindSC aligned the 10 BC subtypes reasonably well (**Fig. 4g**), although separations in the ATAC modality were not as clean as they were in the RNA modality. In comparison, Seurat and LIGER failed to generate meaningful alignments among the BC subtypes (**Fig. 4h-j** and **Supplementary Fig. 9**) while Harmony aligned a few subtypes successfully. These were due partly to the fact that these methods used the low precision gene-level chromatin accessibility profiles as the input (**Fig. 4c**).

Overall, our study demonstrated the power of multiomics in delineating rare cell types and proves that bindSC can generate *in silico* multiomics profiles that are considerably more accurate than do existing tools.

### Integrating scRNA-seq data with spatial transcriptomics (ST) data

BindSC can integrate scRNA-seq data with spatial transcriptomics data to 1) assign spatial locations to cells in the scRNA-seq data and 2) associate additional RNA features to the spatial data for higher resolution delineation. For demonstration, we applied bindSC to integrate the SMART-Seq2 data with the *in situ* spatial transcriptomics data generated by 10x Visium from the same mouse frontal cortex tissue. These two datasets differ widely in number of cells: 1,072 spots in the ST data versus 14,249 cells in the scRNA-seq data (**Supplementary Fig. 10a**). The spots on the Visium assay are at ~50 um resolution and each spot can contain tens of cells. There were 6 clusters identified from the ST data alone, which linked to distinct layers in the corresponding histology images (**Supplementary Fig. 10b-c**) and 23 cell types from the scRNA-seq data alone (**Supplementary Fig. 10d**).

We used bindSC and other programs to derive co-embeddings containing datapoints from both datasets (**Fig. 5a**). BindSC achieved evidently higher alignment mixing scores than the other programs (**Supplementary Fig. 11c**) while the Silhouette scores were similar (**Supplementary Fig. 11b**). For each pseudo-cell in the scRNA-seq data, we calculated its probability to map to a spatial location in the histology image. We then overlaid these cells on the histology image coloring by their probability scores (**Methods**). Noticeably, several cell types in the scRNA-seq data mapped to distinct spatial layers in the histology image, which is consistent with the known cellular anatomy of mouse cortex, particularly for the laminar excitatory neuron cell types such as L2.3 IT, L4, L5.IT, L5.PT, L6.IT, L6.CT, L6B and NP (**Fig. 5b**). Consistent with previous observations, the oligodendrocyte-rich white matter (oligo cells) was mapped below the cortex. BindSC and Seurat were also able to map inhibitory clear cell types such as Lamp5, Vip, Pvalb and Sst in the scRNA-seq data to the histology image, but these cell types did not form distinct spatial patterns. LIGER and Harmony, which had worse alignment mixing scores (**Supplementary Fig. 11c**), failed to map these cells (**Supplementary Figs. 13-14**), especially the Vip cells. The poor mapping of the inhibitory cells may also be attributable to the limited resolution of the Visium technology.

**Fig. 5.**
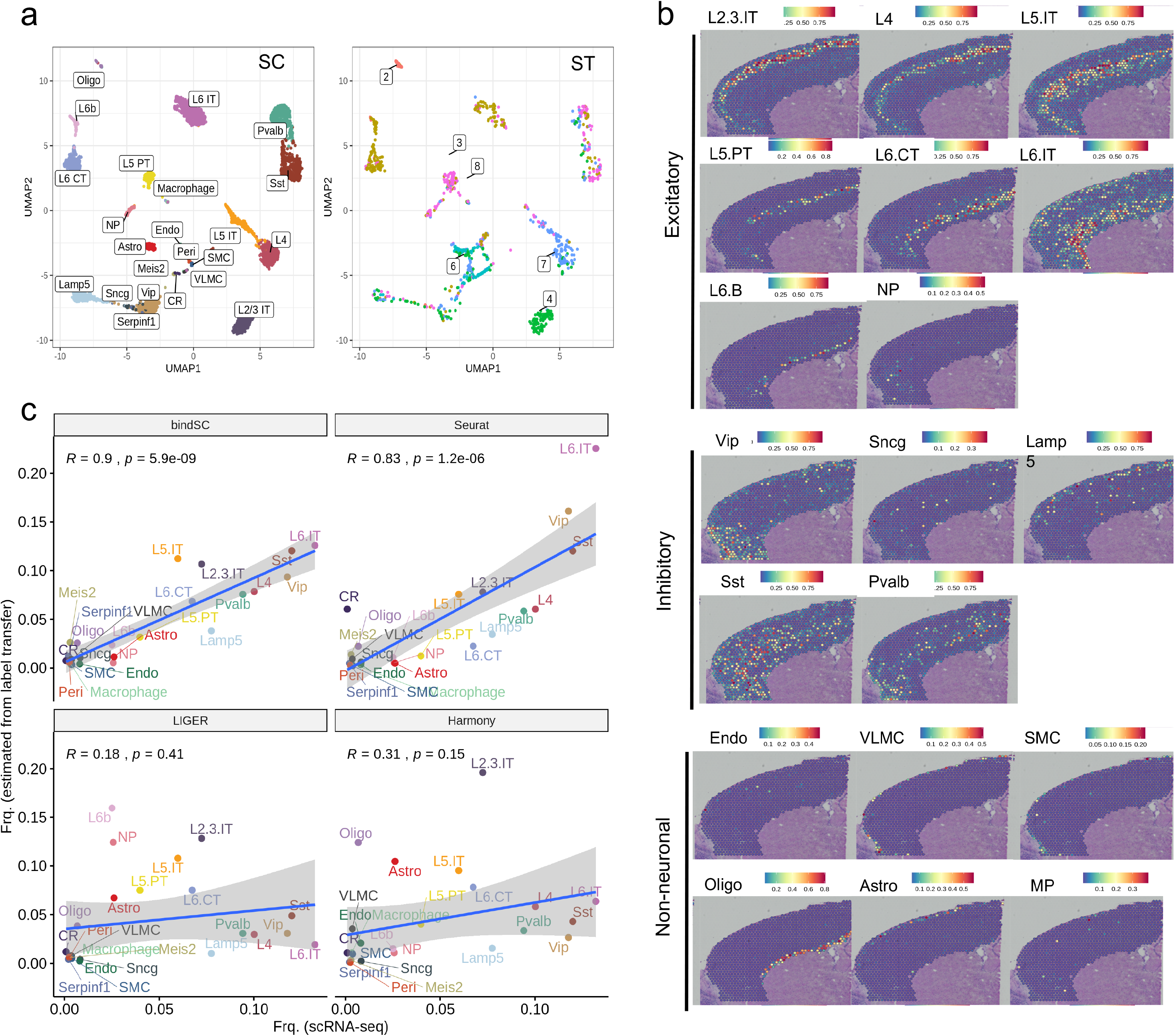
Integrating spatially resolved transcriptomic (ST) data with scRNA-seq data from mouse frontal cortex. UMAPs of the gene expression levels for the 14,249 cells profiled by SMART-Seq2 and for the 1,072 spots profiled by the 10x Visium technology **(a)**. Cell type labels are from the original publication^57^. Predicted locations of each cell type in the histological images **(b)**. Color gradient corresponds to the probability score of a cell being mapped to a particular spatial location. Comparison of cell type frequencies estimated from the ST data (Y-axis) to those estimated from the scRNA-seq data (X-axis) **(c)**. Correlation coefficients (R) and P values are calculated based on Pearson’s correlation analysis. Each dot corresponds to a cell-type (labeled in different colors). The blue line and the grey shade represent regression lines and 95% confidence intervals from performing linear regressions.

Given that each spot in the ST data may encompass multiple cells from multiple cell types, we hypothesized that the probability scores calculated from the co-embeddings can reveal the composition of the cell types at each spot. **Fig. 5c** showed the relationship between cell type abundance in the scRNA-seq data and abundance estimated based on probability scores in the ST data. Results from bindSC achieved the best correlation (Pearson’s R = 0.9). L6.IT, Sst and Vip cell types were the top 3 most abundant cell types in both the scRNA-seq data and the ST data. Seurat also performed reasonably well (Pearson’s R = 0.83) while LIGER and Harmony performed worse. Note that Lamp5 was the cell type that showed the largest discordance in the bindSC result. In examining the spatial distributions of Lamp5 specific gene expressions such as *Lsp1, Npy2r*, and *Dock5,* we could not find any spatial patterns (**Supplementary Fig. 11 d-e**). This finding may indicate that Lamp5 does not have a characteristic spatial distribution.

### Integrating single-cell RNA with protein data

Complex interplay exists between mRNAs and proteins ^37^. Single-cell proteomic methods such as mass cytometry (CyTOF) ^2,38^ measure abundance of a small set of (often 10-50) surface proteins (epitopes) and provide functional quantification of various cell populations. Integrating single-cell RNA and protein data from the same sample can potentially achieve higher resolution characterization and enable discovery of novel cellular states and associated features. BindSC can be applied for such a task. Notice that this task cannot be achieved by any of the existing tools because the mRNA and protein expression levels derived from the same genes are not well correlated, due to complex post-transcriptional modifications and technological limitations ^39^. CITE-seq ^40^ performs jointly profiling of epitope and mRNA levels in the same cells and can be used to evaluate the results of *in silico* integration.

We used a CITE-seq dataset consisting of 30,672 human bone marrow cells with a panel of 25 antibodies ^17^. We split the data into an RNA matrix and a protein matrix. Unsupervised clustering of the RNA matrix revealed cell types largely consistent with those in the protein matrix, except for some noticeable differences (**Fig. 6a-b**). CD8+ and CD4+ T cells were partly blended together in the RNA data but separated clearly in the protein data. On the other hand, conventional dendritic cells (cDC2) were separated from other clusters in the RNA data but were intermixed with other cell types in the protein data. In contrast, unsupervised clustering of the gene expression levels of the 25 protein-homologous RNAs could not yield meaningful classification (**Fig. 6c**). Consequently, Seurat, LIGER and Harmony, which work with only data matrix of 25 homologous features, failed to produce meaningful co-embeddings (**Supplementary Fig. 15**): the cells from the protein data were well clustered, but those from the RNA data were not meaningfully distributed in the co-embeddings.

**Fig. 6.**
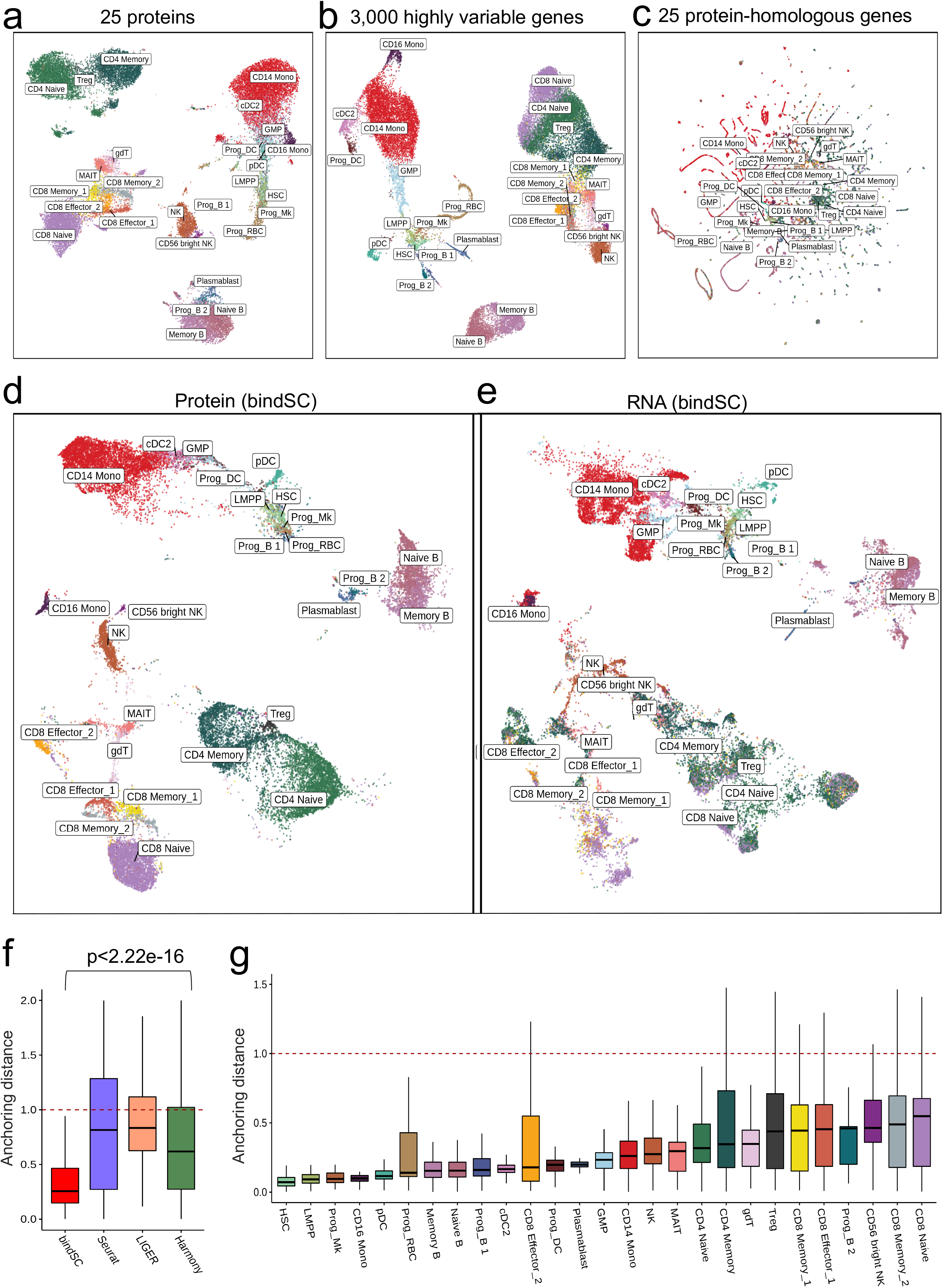
Integrating single-cell RNA with protein data produced by a CITE-seq assay. The UMAP of 30,672 human bone marrow cells based on 25 surface protein levels **(a)**, 3,000 highly variable gene expression levels **(b)** and 25 protein-homologous gene expression levels **(c)**. The cell type labels are from the original study ^17^. UMAP of the protein (**d**) and the RNA (**e**) expression data in the co-embedding generated by bindSC. Comparison of anchoring distances generated by bindSC, Seurat, LIGER and Harmony **(f)**. The red dotted line denotes the anchoring distance from random guesses. Anchoring distances for each cell type in the bindSC co-embedding **(g)**.

We then tested bindSC on this task. The matrix **X** was set as the protein matrix, **Y** the RNA matrix of 3,000 highly variable genes, and **Z** the RNA matrix containing only the 25 protein-homologous genes. Remarkably, the majority of the cells from the two modalities became well aligned in the co-embedding (**Fig. 6d-e**), as they are expected to be. Similar to our previous experiments, we calculated the anchoring distance between the protein and the RNA cells deriving from the same original cells in the co-embeddings. The overall anchoring distance for bindSC was significantly lower than those obtained by Seurat, LIGER, Harmony, or random guesses (p-value < 2.2e-16; Student t-test; **Fig. 6f**). Notably, the bulk of CD4+ and CD8+ T cells in the RNA data became well separated in the co-embedding (**Fig. 6d-e**), thanks to the power of integration. Moreover, the anchoring distances revealed the extent of differences between the levels of the RNAs and those of the homologous proteins in individual cell types (**Fig. 6g**). Interestingly, relatively rare cell types such as HSC, Prog/NK, LMPP, and CD16+ Mono appeared relatively well anchored, whereas relatively common cell types such as CD8 naïve, CD8 memory, B progenitor, Treg, etc. appeared less well anchored. This indicates that there are higher degrees of post-transcriptional heterogeneity in cell types conducting adaptive immune surveillance ^41^.

## Discussion

Despite the ground-breaking advances in single-cell technologies, including multiomics technologies, there always exists a need to computationally integrate multiple data matrices of different modalities from the same biological samples to derive a more comprehensive characterization of cellular identities and functions.

Our method bi-CCA and tool bindSC appeared to have addressed this important analytical challenge without compromising biological complexity in the data. In our experiments, bindSC successfully integrated data obtained from a wide variety of vastly different technologies covering transcriptomes, epigenomes, spatial-transcriptomes and proteomes, and clearly outperformed existing tools such as Seurat, LIGER and Harmony, when being evaluated objectively using true single-cell multiomics data derived from the same cells. In particular, Seurat, LIGER, and Harmony are essentially first-order solutions that can be applied to only rows or columns but not both simultaneously. That approach introduced biases in the results and restricted the utility of those tools in discovering complex cell-type relations and molecular interactions. For instance, they consider only the basic cis-regulatory relations and ignores trans-regulatory relations ^6^ established via distal enhancers, as exemplified in the interaction between *CFLAR* and a 200 kbps upstream putative enhancer site discovered by bindSC and validated by Hi-C in the DEX-treated A549 data. Other scATAC-seq analysis pipelines such as MAESTRO ^16^ and ArchR ^42^ have similar restrictions.

Similarly, bindSC was able to meaningfully associate the expression levels of mRNAs with those of the surface proteins, a very challenging task due to complexity in post-transcriptional modification. The resulting co-embedding offered deeper biological insights than embeddings derived from single modality or by using other existing approaches. For example, CD4+ T cells became evidently separated from CD8+ T cells and so did pDC cells from other cell types.

BindSC also achieved meaningful mapping of scRNA-seq data to spatial locations in the brain cortex samples, after integrating with the ST data. Even though the two datasets were not both at single-cell resolution, bindSC was still able to achieve a meaningful integration.

Bi-CCA made two assumptions: 1) the two sets of cells are sampled uniformly from the same biological sample; 2) the features of the two datasets are linearly correlated. These two assumptions are met under many scenarios of current investigations, however, could be violated when there are insufficient number of cells obtained from a rapidly developing cell population. Consequently, the accuracy of the co-embedding could vary, depending on the sampling density and the complexity of the population. We measured accuracy with respect to data complexity in the simulation experiments, however, accuracy on a real dataset could be complex to gauge *a priori* and will require case by case investigation in the context of a specific study, followed by necessary experimental validation. Nonetheless, in this study we clearly proved based on objective ground truth data that bi-CCA substantially avoided bias introduced by existing methods and that bindSC is a robust implementation that can be applied to derive meaningful results on most recent datasets containing thousands to tens of thousands cells (**Supplementary Table 1**).

BindSC is efficiently implemented in R. The major computational cost for bindSC is from calculating cell/feature co-embedding coordinates using singular value decomposition (SVD) (**Methods**); It typically requires *O(MNL)* floating-point operations to construct *MN* cell-cell distance matrix as input to SVD decomposition, where *M* and *N* are cell number of the two modalities, respectively. To address this computational challenge, bindSC implements the “divide-and-conquer eigenvalue algorithm”. The divide part first splits cells into different blocks specified by users, which can be solved in parallel with lower memory usage (**Supplementary Fig. 1b**). The conquer part then merges results from each block recursively. Therefore, the maximal memory usage of bindSC is independent of the total cell number.

Taken together, we believe that bindSC is likely the first tool that has achieved unbiased integration of data matrices generated by different technologies and can be applied in broad settings. In the single-cell domain, bindSC can clearly be applied to align cells and features simultaneously, which are important for ongoing investigations in the Human Cell Atlas ^43^, the NIH HubMap ^44^, the Human Tumor Cell Network ^45^ and on remodeling of tumor microenvironment ^46^. Further, bindSC can potentially be applied to other domains, such as integrating patient sample mRNA profiles with cell-line drug-sensitivity data ^47^.

## Methods

### BindSC workflow

BindSC workflow for creating *in silico* single cell multi-omics embeddings consists of five steps:

1. individual dataset preprocessing including variable feature selection and cell clustering,
2. initializing feature matching across modalities (i.e., constructing gene score matrix),
3. identifying cell correspondence using the bi-CCA algorithm,
4. jointly clustering cells between two modalities in the co-embedding latent space and constructing pseudo-cell level multi-omics profiles, and
5. downstream analysis for various integration tasks.

We formulate our method for the case of two modalities. Let 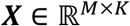 be a single-cell dataset of features *g_1_,g_1_,…,g_M_* by cells *c_1_,c_1_,…,c_K_* and 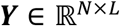 be a single-cell dataset of feature *P_1_,P_2_,…,P_N_* by cells *d_1_, d_1_,…, d_L_*. *M* and *N* are the numbers of features (e.g., gene expression, chromatin accessibility, protein abundance level) in the two datasets. *K* and *L* are the number of cells in the two datasets. Without loss of generality, we assume that features *g_1_, g_1_,…, g_M_*represent the gene expression levels and *M* ≤ *N*. The important component of each step is described as follows.

#### 1. Individual modality preprocessing

For each modality, we follow standard processing pipeline, which includes variable feature selection and unsupervised cell clustering. The cluster information derived from each modality is used for downstream parameter optimization.

#### 2. Initializing feature matching across modalities

Because features in the two datasets are generally different, bindSC requires one additional transition matrix 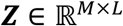 as input for bridging the integration of ***X*** and ***Y***. The transition matrix ***Z*** can be understood as the projection of ***Y*** to the feature space of the first dataset ***X***. Taking the integration of scRNA-seq and scATAC-seq as an example, the matrix Z can be derived from scATAC-seq profiles by summing reads in gene bodies ^17,19,23^. This can also be input from the regulatory potential (RP) model in MAESTRO ^16^. In a simpler case where ***X*** and ***Y*** have matched features, the integration tasks fall into two categories: 1) batch correction for scRNA-seq data across individuals, species, or technologies; 2) integration of scRNA-seq with spatial transcriptome data. In those cases, the transition matrix ***Z*** is initialized as ***Y***. In bi-CCA, ***Z*** is updated iteratively. In the following text, the initial value of ***Z*** is denoted by ***Z***^0^.

#### 3. Bi-order canonical correlation analysis (Bi-CCA)

The key algorithm implemented in bindSC is Bi-CCA, the concept of which extends traditional CCA^17,24,48^ to both rows and columns to enable capturing of correlated variables in cells and features simultaneously. Bi-CCA introduces two cell-level projection matrices 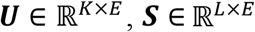 such that the correlations between indices ***XU*** and ***ZS*** are maximized, and two feature-level projection matrices 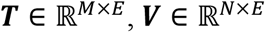 such that the correlations between indices ***Z’T*** and ***Y’V*** are maximized. The optimization framework can be formulated as:

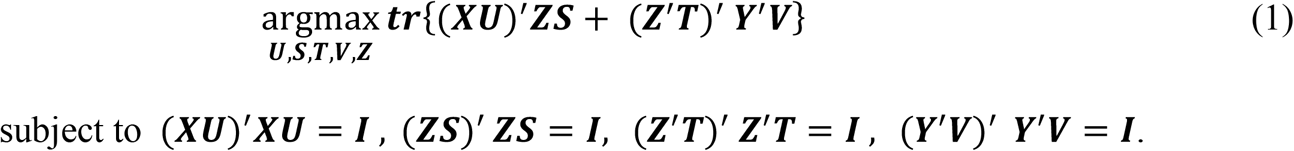

If the transition matrix ***Z*** is known, the objective (1) can be divided into two disjoint traditional canonical correlation analysis (CCA) problems. The left term is performed to identify cells of similar (aligned) features, while the right term is performed to identify features shared by the (aligned) cells, each of which can be solved in the CCA framework. However, it is difficult to update transition matrix ***Z*** in equation (1) even when matrices ***U,S,T,V*** are available. This is because: a) left optimization problem requires ***Z*** as input and the right optimization problem requires ***Z’*** as input, leading (1) to a non-linear optimization problem; b) transition matrix ***Z*** shows up in constraints.

Therefore, we modify equation (1) in a much more practical way. First, we standardize ***X*** to let it have ***X’X = I***, and standardize ***Y*** so that ***’ = I***. The standardization process can be seen in **Algorithm 1**. Thus, equation (1) could be simplified as

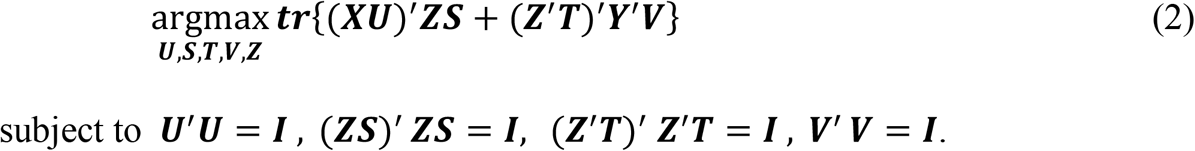

To eliminate transition matrix ***Z*** from constraints, we introduce two transition matrices 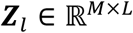 and 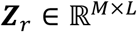 and optimize the following problem:

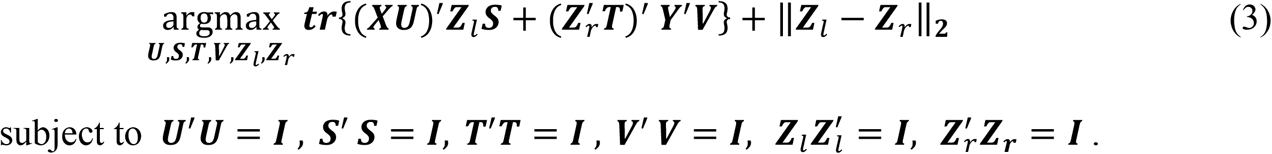

To solve equation (3), we also standardize ***Z^(0)^*** to let ***Z^(0)^ Z^(0)^ = I***, and initialized with *Z_l_: = Z^(0)^*. The standard singular value decomposition (SVD) can be implemented to obtain the canonical correlation vectors (CCVs) at cell levels. We used a user-defined number (E) of singular vectors that approximate the CCVs (**Algorithm 2**). Here we term *E* to represent the cell-level “dimensionality” in the latent space, which is a parameter required to be optimized (Details seen in **Parameter optimization**).

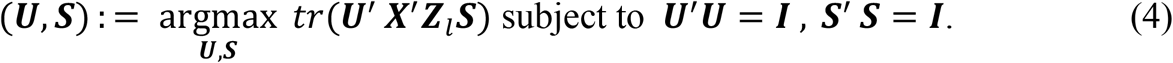

Having CCV pair (***U, S***) obtained, we have cell correspondence in the latent space between two datasets. The left transition matrix ***Z**_l_* can be updated by:

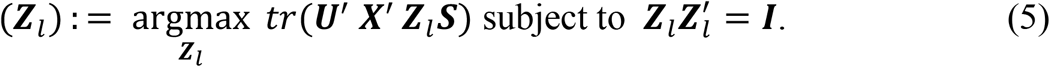

The details of solving optimization problem (5) is in **Algorithm 2**.

We then set

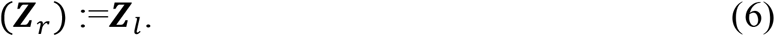

The similar SVD algorithm (**Algorithm 2**) is used to approximate CCVs:

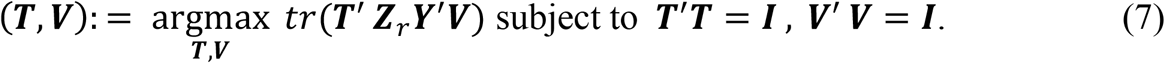

Once CCV pairs (***T, V***) are obtained, the features are matched in the latent space between two datasets. The right transition matrix ***Z**_r_* could be updated as:

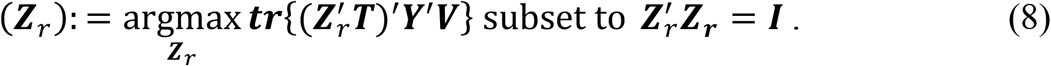

Next, we set

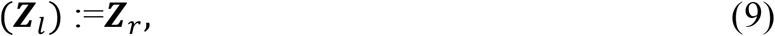

The update process (4) ~ (9) are repeated until convergence. Because each of the subproblems is convex with respect to the block variables being optimized, the algorithm is guaranteed to converge to a fixed point (local minimum).

In the above framework, the transition matrix ***Z*** (represented by ***Z**_l_* and ***Z**_r_*) is updated based on original observed matrices ***X*** and ***Y***. In practice, we introduce the couple coefficient *α* (0≤ *α ≤* 1) to assign weights on initialized matrix ***Z^(0)^*** on transition process (6) and (9).

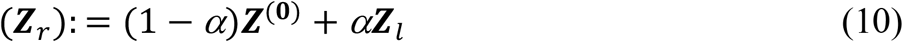

and

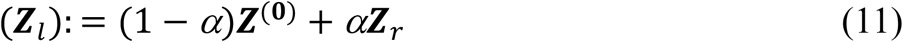

The couple coefficient *α* can reflect the contribution of initial ***Z^(0)^*** on linking two modalities. Equations (10) and (11) will be reduced to Equations (6) and (9) if *α* = 1. The bi-CCA algorithm will be reduced to traditional CCA if *α* = 0. Selection of coefficient *α* can be seen in **Parameter optimization**. Notably, the final ***Z**_r_* and ***Z**_l_* will be converged to different matrices if *α* < 1. The workflow of the iterative process is shown in **Supplementary Fig. 1a**.

### Jointly clustering cells across datasets in shared latent space and constructing pseudo-cell level multi-omics profiles

Equation (4) projects cells of two datasets into a correlated *E*-dimensional space with cell coordinates ***U = (u_1_,u_2_,…,u_K_***) and ***S = (s_1_,s_2_,…,s_L_***), respectively. L2-normalization is performed to remove global differences in scale, therefore

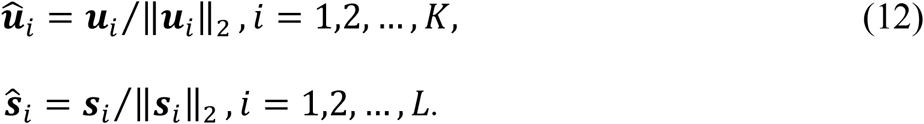

The shared nearest neighbor (SNN) graph is constructed by calculating the *l*-nearest neighbors (20 by default) based on the Euclidean distance of L2-normlized latent space. The fraction of shared nearest neighbors between the cell and its neighbors is used as weights of the SNN graph. The modularity optimization technique Leiden algorithm ^49^ is used to group cells into interconnected clusters (termed meta-cluster) based on constructed SNN graph with a resolution parameter setting by users (default 0.5).

To understand the molecular-level interaction among modalities, we construct the pseudo-cell level multi-omics profiles. Briefly, for cells in each meta-cluster identified, the Leiden algorithm is further performed based on SNN graph with a higher resolution (default = 2). In this way, cells in each meta-cluster are further grouped into highly interconnected sub-clusters. We term such sub-clusters as pseudo-cells. Only pseudo-cells that consist of at least *n* cells (default = 10) are kept for downstream analysis, while the others are considered data-specific and discarded. Profiles of the pseudo-cells are constructed by aggregating the cells included. We denote by 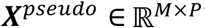 be pseudo-cell profiles of feature *g_1_, g_1_,…,g_M_* and 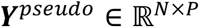 be pseudo-cell profiles of feature *p_1_,p_2_,…,p_N_*. *P* is the number of pseudo-cells.

### Algorithm 1. Standardizing inputs

For input matrix *X*, we denote 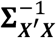 as the generalized inverse of matrix ***X’X***, and redefine 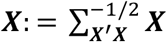. For input matrix ***Y***, we denote 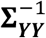 as the generalized inverse of matrix ***YY’***, and redefine 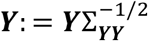. The standardization of ***Z**_r_* and ***Z**_l_* is the same as above.

### Algorithm 2. Calculating CCVs using SVD

Take subproblem from the Equation (4) as an example, the goal of this module is to find projection matrix 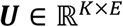 and 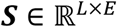 such that the correlations between two indices ***XU*** and ***Z**_l_**S*** are maximized.

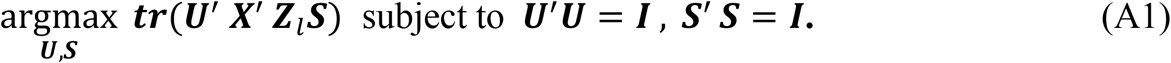

We define 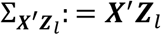. Let 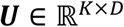 and 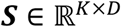 be the matrices of the first *E* left- and right singular vectors of 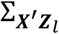. Then the optimum in Equation (A1) is solved.

### Algorithm 3. Updating transition matrix with orthogonality constraints

Take subproblem from the Equation (5) as an example, the goal of this module is to optimize ***Z**_l_*.

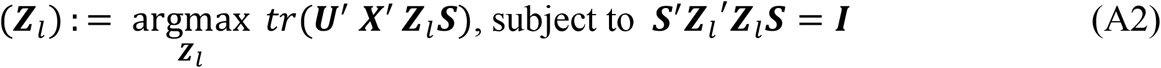

Equation (A2) is maximized when ***Z_l_S = XU***. Therefore, we can update ***Z_l_*** as

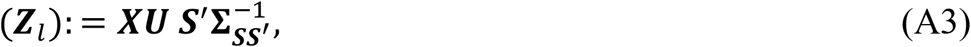

where 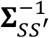 denotes the generalized inverse of matrix ***SS’***.

### Parameter optimization

There are two key hyperparameters when running bindSC for integration. The first one is the dimensionality *E* in the latent space and the second one is the couple coefficient *α*. Similar with previous integration methods, the number of dimension *E* is very important on cell type classification. We provide heuristics to guide the selection of *E* based on integration metrics defined below, though sometimes helpful, are not substitute for biological insights. As a general suggestion, we recommend starting with a value of *E* the same with the minimal number of principle components (PCs) used on single modality clustering. The selection of couple coefficient *α* depends on whether initialized *Z^(0)^* can represent the gene score of *Y*. We devise two metrics to aid in selecting *α*, which measure integration performance on accuracy (no mixing of cell type) and alignment (mixing of datasets) as defined below.

#### 1) Silhouette score

To measure integration accuracy, we use the Silhouette score. Cluster for each cell is defined using the cell type labels assigned from single dataset clustering. The Silhouette score assesses the separation of cell types, where a high score suggests that cells of the same cell type are close together and far from cells of a different type. The Silhouette score *s*(*i*) for each cell is calculated as following. Let *a*(*i*) be the average distance of cell *i* to all other cells within i’s cluster and *b*(*i*) the average distance of *i* to all cells in the nearest cluster, to which cell *i* does not belong. Cell-cell distance is computed in the L2-normalized co-embeddings (**Equation 12**). *s*(*i*) can be computed as:

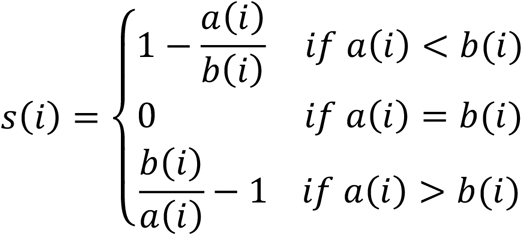

Notably, given accurate correspondence between two modalities unknown, calculating *s*(*i*) for cell *i* in above equation only includes cells from the same dataset. We average values across all cells to obtain an overall silhouette score for integration task.

#### 2) Alignment mixing score

To measure integration mixing level, we use an alignment mixing score similar to those of previous studies ^50^. We first build a 20-nearest neighbor graph for each cell from L2-normalized co-embeddings (**Equation 12**). For cell *i*, assuming proportions of cells from two modalities are *p_1i_* and *p_2i_*, respectively, the alignment mixing score is calculated as

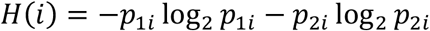

This corresponds to a mixing metric per cell, and we average values across all cells to obtain an overall mixing metric.

We run bindSC by ranging a from 0 to 1 (with step size 0.1). Silhouette score and alignment mixing score is calculated for each scenario. We select appropriate a that generally has best performance in Silhouette score and alignment mixing score. Parameter values used in this study can be seen in **Supplementary Table S1**.

### Performance and benchmarking

In our evaluation, in addition to Silhouette score and alignment mixing score, we also consider anchoring distance for evaluation datasets from multi-omics technologies, in which each cell has paired profiles. For cell *i* from the first data, we calculate its distance (Euclidean distance) with all cells in the second data as *D_i_*, and its distance with cell *i* in the second data as *d_i_*. The anchoring distance for cell *i* is calculated as *2d_i_/max(D_i_*). We then average anchoring distance across all cells to obtain an overall anchor distance metric. The anchoring distance of cell *i* is 0 when it is anchored correctly. The overall anchoring distance is 1 if we randomly layout cells on co-embeddings.

### Simulation dataset

We generated simulation dataset to evaluate method performance in integrating two modalities assuming accurate feature matching unknown. We used Splatter tool ^26^ to simulate dataset *X* with 1,000 genes and cells with different sizes (from 1,000 to 10,000). The whole population is consisted of three batches (cell types). To mimic the feature unmatching case, we first generated dataset *X_1_* by randomly permutating genes of *X* under specified misalignment rates (MR). MR ranges from 0 to 1 with step size being 0.05 in this study. *X_1_* is the same as *X* if MR = 0. Half of genes are matched between *X_1_* and *X* if MR = 0.5. No genes are matched between *X_1_* and *X* if MR = 1. Then we generated matrix *Z* by adding *X_1_* with white noise at certain level (i.e., Signal-Noise-Ratio; SNR). SNR is set to be three levels (0, 0.25 and 0.5).

For method comparison, previous methods including traditional CCA, Seurat ^17^, Liger ^19^, and Harmony tools ^18^ take *X* and *Z* as input assuming that cell correspondence between them is unknown. bindSC takes two parts as input: 1) *X* and *Z* with cell correspondence unknown; 2) *X* and *Z* with feature-level matching unknown (**Supplementary Fig. 2**).

### Preparation of dexamethasone (DEX) treated A549 cell dataset

To investigate the ability of bindSC in integrating scRNA-seq and scATAC-seq profiles, we explored the DEX-treated A549 dataset generated from sci-CAR technology, which uses combinatorial indexing-based assay to jointly profile chromatin accessibility and mRNA on same cell ^6^. In the A549 dataset, DEX is a synthetic corticosteroid which activates glucocorticoid receptor (GR), binds to thousands of locations, and alternates the expression of hundreds of genes ^51^. The human lung adenocarcinoma derived A549 cells after 0, 1, or 3 hours of 100nM DEX treatment are assayed. The sci-RNA-seq dataset was from GSE117089 (https://www.ncbi.nlm.nih.gov/geo/query/acc.cgi?acc=GSE117089) and sci-ATAC-seq data was from GSM3271041 (https://www.ncbi.nlm.nih.gov/geo/query/acc.cgi?acc=GSM3271041). The original A549 data includes sci-RNA-seq profiles for 6,150 cells and sci-ATAC-seq profiles for 6,260 cells. There are 1,429 cells co-assayed. Following Cao et al., pre-processing pipeline (https://github.com/KChen-lab/bindSC/blob/master/vignettes/A549/A549_preprocess.ATAC.Rmd), we binarized peak count matrix for cells from both ATAC-seq only and co-assay. Loci present in less than 5 cells and cells with less than 300 accessible loci were removed. Peaks within 1kb were merged and reads in merged peaks were aggregated to generate a merged peak matrix, leading to 3,628 cells with 32,791 loci. Each locus’ accessibility in each cell was calculated by dividing the cell’s raw read count by cell specific size factor using *estimateSizeFactors* function in *Monocle 2* ^52^. For RNA-seq data, cells with expression counts less than 500 and more than 9100 were removed. The gene expression in each cell was also calculated by dividing the cell’s raw read count by cell specific size factor, followed by *log2* normalization. Genes with no variation in expression across cells were further removed. The gene activity matrix was collapsed from the peak matrix by summing all counts with the gene body plus 2kb upstream using *CreateGeneActivityMatrix* function in Seurat3 ^17^. We then picked top 10,000 variable genes in both sci-RNA-seq data and gene activity data and used the overlapped 4,759 genes between them for integration. Finally, the sci-RNA-seq matrix was composed of 6,005 cells with 4,759 genes, the gene activity matrix was composed of 3,628 cells with 4,758 genes, and the sci-ATAC-seq matrix was composed of 3,628 cells with 24,953 loci. There were 1,429 cells co-assayed.

### Preparation of the mouse skin cell data

We examined the performance of bindSC in integrating the scRNA-seq and scATAC-seq data derived from mouse skin tissue. This dataset was generated using SHARE-seq (3) which included 34,774 cells that have joint profiles of RNA and ATAC profiles. The RNA data was downloaded from https://www.ncbi.nlm.nih.gov/geo/query/acc.cgi?acc=GSM4156608. The ATAC data was downloaded from https://www.ncbi.nlm.nih.gov/geo/query/acc.cgi?acc=GSM4156597. The final ATAC-seq matrix includes 25,594 cells on 74,161 peaks after quality control (including removing cells with less than 350 genes expressed; peaks that exist in less than 500 cells). In addition, 4,894 genes were identified that were highly variable in both gene expression and gene activity profiles. For this evaluation, we only focused on the third metric (e.g., anchoring distance) that represents the chance for the two instances of a co-assayed cell to appear in the co-embeddings.

### Preparation of the mouse retina 10x Genomics Multiome ATAC+RNA data

One mouse retina was dissociated by papain-based enzymatic digestion as described previously ^53^ with slight modifications. Briefly, 45 U of activated papain solution (with 1.2 mg L-cysteine (Sigma) and 1200U of DNase I (Affymetrix) in 5ml of HBSS buffer) was added to the tissue and incubated at 37 °C for 20 minutes to release live cells. Post-incubation, papain solution was replaced and deactivated with ovomucoid solution (15 mg ovomucoid (Worthington biochemical) and 15 mg BSA (Thermo Fisher Scientific) in 10 ml of MEM (Thermo Fisher Scientific)). The remaining tissue clumps were further triturated in the ovomucoid solution and filtered through a 20nm nylon mesh. After centrifugation at 300g 10min at 4C, the singe cells were resuspended PBS with 0.04% BSA and checked for viability and cell count. About 1 million cells were pelleted and resuspend in chilled lysis buffer (10x Genomics), incubate for 2 minutes on ice while monitored under microscope. 1ml of chilled wash buffer (10x Genomics) was added and sample was spun down at 500g 5min at 4C and washed before resuspended in Diluted Nuclei Buffer (10x Genomics). Nuclei concentration was determined using countess and proceed with transposition according to manufacturer’s recommendation (10x Genomics). After incubation for one hour at 37C, the transposed nuclei were combined with barcoded gel beads, RT mix and partition oil on Chromium to generate gel beads in Emulsion (GEMs). Single cell ATACseq library and 3’RNAseq library were subsequently generated following recommended protocol from 10x Genomics. Libraries were quantified and loaded on Novaseq 6000 and run with the following parameter: 151, 8, 8, 151bp. Data was analyzed using bcl2fastq (to generate fastq files) and cellranger pipeline (10x Genomics).

### Preparation of the mouse frontal cortex cell data

We investigate bindSC ability in integrating spatially resolved transcriptomic (ST) with dissociated scRNA-seq. For the ST dataset, we used sagittal mouse brain slices generated from the Visium v1 chemistry. The dataset was downloaded from https://support.10xgenomics.com/spatial-gene-expression/datasets. The pre-processing workflow was guided by the Seurat3 (https://satijalab.org/seurat/v3.2/spatial_vignette.html). Briefly, cells were subset from anterior region, followed by *sctransform* ^54^. We then proceed to run dimensionality reduction and clustering using standard workflow as did for scRNA-seq. Cluster ID 1,2,3,5,6,7 was extracted, followed by segment based on exact position (Details in **Subset out anatomical regions** part in Seurat3 tutorial), leading to 1,072 cortical cells left for the ST data. One cortical scRNA-seq data composed of ~14,000 adult mouse cortical cell taxonomy from the Allen Institute was collected (https://www.dropbox.com/s/cuowvm4vrf65pvq/allen_cortex.rds?dl=1). This dataset was generated using the SMART-Seq2 protocol ^55^. The *sctransform* normalization was performed based on 3,000 variable genes. We used the cell type annotation provided by published meta data available. There was a total of 14,294 cortical cells with 34,617 genes for the scRNA-seq data. Integration of scRNA-seq and ST is based on 2,316 variable genes overlapped between two datasets.

To predict locations of each cell type from scRNA-seq in the histological images, we built a support vector machine (SVM) that trained on cell profiles from scRNA-seq data. In the training model, features were identified as cell coordinates in co-embeddings and labels were corresponding cell types. The trained SVM was applied to ST data and output predicted probability of each cell type at each spot. The *SpatialFeaturePlot* function in Seurat3 was used to overlay predicted probabilities for each cell type on top of tissue histology.

### Preparation of human bone marrow cell dataset

We examined the performance of bindSC in integrating the single-cell RNA and protein data derived from human bone marrow tissue. This dataset was generated using the CITE-seq technology ^40^, which included 30,672 cells that have joint profiles of RNA and a panel of 25 antibodies. The dataset was downloaded from https://satijalab.org/seurat/v4.0/weighted_nearest_neighbor_analysis.html. We extracted the 25 protein-homologous gene expression profile from the RNA data and kept cells that have total expression count > 2. The final protein matrix includes 28,609 cells with 25 protein abundance levels. The gene expression matrix includes 28,609 cells with 3,000 genes. The protein-homologous RNA matrix includes 28,609 cells with the RNA levels of the 25 genes homologous to the 25 proteins. To measure anchoring accuracy for each cell type, we used the third metric, anchoring distance, which measures the distance of protein and gene expression for each cell in co-embeddings.

### Motif-based Transcription Factors (TFs) activity estimation

To estimate transcription factor activity from scATAC-seq data, we used default settings in chromVAR ^56^ package. This approach quantifies accessibility variation across single cells by aggregating accessible regions containing a specific TF motif. It calculated motif-based TF activity by comparing the observed accessibility of all the peaks containing a TF motif to a background set of peaks normalizing against known technical confounders.

## Acknowledgements

This project has been made possible in part by the Human Cell Atlas Seed Network Grant CZF2019-02425 to RC and KC, CZF2019-002432 to KC from the Chan Zuckerberg Initiative DAF, an advised fund of Silicon Valley Community Foundation, grant R01EY022356 and R01EY018571 to RC from National Eye Institute, grant RP180248 to KC from Cancer Prevention & Research Institute of Texas, grant U01CA247760 to KC and the Cancer Center Support Grant P30 CA016672 to PP from National Cancer Institute. This project was also partially supported by the Single Cell Genomics Core at Baylor College of Medicine funded by NIH shared instrument grants (S10OD023469, S10OD025240) and P30EY002520. The authors would like to thank Qingnan Liang, Yuanxin Wang, Linghua Wang, Tapsi Kumar, Runmin Wei, Nicholas Navin, John Weinstein and Hussein Abbas for their comments.

## Author contributions

K.C. conceptualized and supervised the project. J.D. designed the bindSC tool, implemented the software and performed analysis. R.C., Y. L., X. C., S.K., J.C., contributed to mouse retina 10x Genomics ATAC+RNA data generation, curation. V.M. contributed to data interpretation. J.D., S.L. and K.C. drafted the manuscript. All authors reviewed, edited, and approved the manuscript.

## Competing interests

The authors declare no competing interests.

## Notes

### Competing Interest Statement

The authors have declared no competing interest.

